# Loss of IKK subunits limits NF-κB signaling in reovirus infected cells

**DOI:** 10.1101/843680

**Authors:** Andrew J. McNamara, Pranav Danthi

## Abstract

Viruses commonly antagonize innate immune pathways that are primarily driven by Nuclear Factor-κB (NF-κB), Interferon Regulatory Factor (IRF) and Signal Transducer and Activator of Transcription (STAT) family of transcription factors. Such a strategy allows viruses to evade immune surveillance and maximize their replication. Using an unbiased RNA-seq based approach to measure gene expression induced by transfected viral genomic RNA (vgRNA) and reovirus infection, we discovered that mammalian reovirus inhibits host cell innate immune signaling. We found that while vgRNA and reovirus infection both induce a similar IRF dependent gene expression program, gene expression driven by the NF-κB family of transcription factors is lower in infected cells. Potent agonists of NF-κB, such as Tumor Necrosis Factor alpha (TNFα) and vgRNA, failed to induce NF-κB dependent gene expression in infected cells. We demonstrate that NF-κB signaling is blocked due to loss of critical members of the Inhibitor of KappaB Kinase (IKK) complex, NF-κB Essential MOdifier (NEMO) and IKKβ. The loss of the IKK complex components prevents nuclear translocation and phosphorylation of NF-κB, thereby preventing gene expression. Our studies demonstrate that reovirus infection selectively blocks NF-κB, likely to counteract its antiviral effects and promote efficient viral replication.

**IMPORTANCE:** Host cells mount a response to curb virus replication in infected cells and prevent infection of neighboring, as yet uninfected cells. The NF-κB family of proteins is important for the cell to mediate this response. In this study, we show that in cells infected with mammalian reovirus, NF-κB is inactive. Further, we demonstrate that NF-κB is rendered inactive because virus infection results in reduced levels of upstream intermediaries (called IKKs) that are needed for NF-κB function. Based on previous evidence that active NF-κB limits reovirus infection, we conclude that inactivating NF-κB is a viral strategy to produce a cellular environment that is favorable for virus replication.

## INTRODUCTION

The mammalian innate immune response is an effective response to viral intrusion. The primary mechanism of innate control of virus infection is the production of antiviral cytokines. Paracrine signaling by these cytokines establishes an antiviral state in neighboring, uninfected cells, making them refractory to virus infection and limiting dissemination of virus in the host (1). Expression of these cytokines is under the control of two major transcription factors, Nuclear Factor-κB (NF-κB) and Interferon Regulatory Factor 3 (IRF3) (2). NF-κB and IRF3 are activated downstream of a signal that is initiated by sensing of pathogen-associated molecules extracellularly, within transit through cellular uptake pathways, or within the cell (2). For RNA viruses, cell surface or endosomal sensing of the genomic material via Toll like Receptors (TLRs), or cytoplasmic sensing via the RIG-I like Receptors (RLRs) are two major mechanisms of pathogen recognition (2). Animal models and cell lines lacking any component of the signaling module – sensor, transcription factors, or cytokines – are typically more susceptible to viral infections than those with an intact immune response (3–6).

Because these initial stages of the innate immune response are so effective at limiting viral replication, most viruses have evolved one or more mechanisms to limit either the production or activity of these anti-viral cytokines (7, 8). Frequently, viruses antagonize the immune response by sequestering, degrading, or inactivating one or more cellular components that are required for a cytokine based antiviral response (7, 8). Targeting transcription factor function is a commonly used viral strategy, perhaps because transcription factors serve a critical node that controls the expression of multiple antiviral molecules. Among these, NF-κB can control the function of a wide variety of pro-inflammatory chemokines and cytokines (9). The NF-κB transcription factor family is composed of five different subunits that function as homo or heterodimers. The classical NF-κB complex (henceforth referred to as NF-κB), composed of p65 and p50 subunits, is a critical regulator of antiviral gene expression (9). In an inactive state, it is sequestered in the cytoplasm by the Inhibitor of κB (IκB) inhibitor proteins (9). NF-κB transcriptional activity is regulated by the IκB Kinase (IKK) complex (9). The IKK complex, which is composed of IKKα, IKKβ, and NEMO, phosphorylates IκB, which leads to its ubiquitination and degradation. Freed from its inhibitor, NF-κB is able to translocate to the nucleus, bind to DNA, and initiate gene expression. The transactivation function of NF-κB also requires IKK-mediated phosphorylation of the p65 subunit (10).

Mammalian orthoreovirus (reovirus) is a dsRNA virus which replicates in the cytoplasm of the host cell (11). Like most other viruses, reovirus pathogenesis is influenced by NF-κB signaling (12). In a newborn mouse model, NF-κB plays an antiviral role in the heart. In comparison to wildtype mice, NF-κB p50 −/− mice exhibit higher viral titers, tissue damage, and cell death, indicating that NF-κB is antiviral in this context. This outcome, at least in part, is due an inability of p50−/− mice to produce IFNβ. Despite the fact that the viral genomic RNA remains within the two concentric protein shells that comprise the reovirus capsid, the current model posits that the innate immune response is initiated when genomic RNA from incoming virions is sensed by the RLRs - RIG-I and MDA5 (9, 13). The sensing of the RNA leads to the activation of IRF3 and NF-κB, which lead to the production of IFN and other inflammatory cytokines (14). Whether reovirus actively limits this antiviral response has not been extensively scrutinized.

In this study, we investigated whether reovirus inhibits innate immune signaling following infection. Using RNA-seq, we found that NF-κB activity was inhibited in infected cells following treatment with multiple agonists including viral genomic RNA (vgRNA) and Tumor Necrosis Factor alpha (TNFα). We discovered that this inhibition was due to reduced cellular levels of the IKK components, IKKβ and NEMO. Loss of the IKK complex led to inhibition of NF-κB nuclear translocation and consequent blockade of its transactivation function. Blockade of viral gene expression prevented IKK loss, suggesting that events in viral replication after cell entry are required for IKK loss and NF-κB inhibition. This study highlights a previously unknown mechanism by which reovirus infection blunts the host innate immune response.

## RESULTS

### NF-κB dependent gene expression is blocked in reovirus-infected cells

The genomic dsRNA within reovirus particles serves as the pathogen associated molecular pattern that activates the innate immune response via RIG-I and MDA5 (15–17). To determine if reovirus infection modifies this response, we compared the host cell response in L929 cells following transfection of vgRNA with the response following infection with reovirus strain type 3 Abney (T3A). Using RNA-seq analyses, we found that viral RNA transfection induced the expression of 978 genes (Fig. 1A, 1B). For these analyses, we considered only those genes whose expression was increased > 4 fold (log_2_FC > 2) and were identified with false discovery rate (FDR) of < 0.05 to be significantly different. We used iRegulon, which predicts transcriptional regulators for a similarly expressed gene set by providing a Normalized Enrichment Score (NES) (18). A high NES for a given transcription factor indicates that many of the genes in a set are likely regulated by that transcription factor. We used this program to identify which transcription factors most likely regulate the genes induced following treatment with vgRNA. We found that, of the 978 genes induced by vgRNA, the highest NES scores were assigned to NF-κB and IRF, with scores of 4.0 and 10.0 respectively (Fig. 1C). Predictably, reovirus infection induced a similar gene expression profile. 65% of the 978 genes induced by vgRNA were also induced by reovirus. When we used iRegulon to predict the transcription factors which regulate the genes induced by reovirus infection, we found that, while genes regulated by IRF were enriched in this list (NES of 12.0), genes regulated by NF-κB were not. Surprisingly, the NES for NF-κB fell below the 3.0 cutoff indicating that NF-κB target genes were not enriched in the set of genes induced by reovirus infection (Fig. 1C). Of the 978 genes induced by vgRNA described above, 35% of genes (339 of 978) were expressed to a lower extent in reovirus infected cells. Using iRegulon, we predicted that NF-κB target genes were enriched in this set, as the NES for NF-κB increased from 4.0 to 4.9 (Fig. 1C). These data suggest that NF-κB and IRF transcription factor families are regulated differently in cells transfected with RNA and cells infected with reovirus. Thus, the observed differences in the gene expression profiles of RNA transfected and reovirus infected cells are not related to differences in RNA sensing. Instead, this difference may be because reovirus fails to activate NF-κB signaling pathway or because it has evolved a mechanism to block NF-κB signaling.

**Fig 1.**
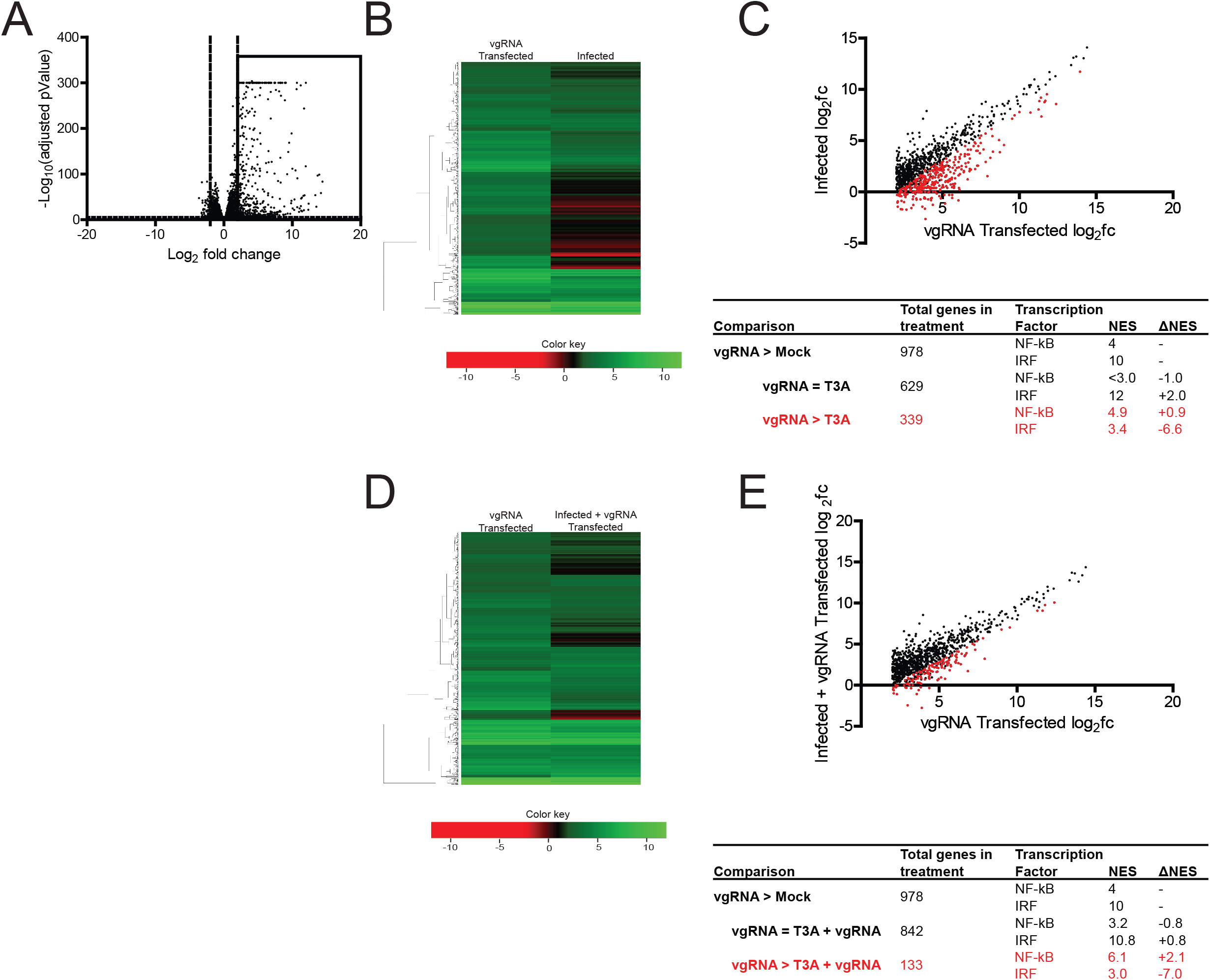
Reovirus strain T3A inhibits NF-κB dependent gene expression. (A) ATCC L929 cells were transfected with 0.5 μg of vgRNA. 7 h following transfection, total RNA was extracted and subjected to RNA-seq analyses. A volcano plot showing genes whose expression is induced > 4-fold (log_2_FC>2) with FDR < 0.05 in comparison to mock infected cells are shown within the box. (B, C) ATCC L929 cells were transfected with 0.5 μg of vgRNA for 7 h or infected with 10 PFU/cell of reovirus strain T3A for 20 h. Total RNA was extracted and subjected to RNA-seq analyses. (B) A heat map comparing expression of genes shown in boxed region of Fig. 1A following vgRNA transfection and T3A infection is shown. (C) A scatterplot comparing expression of genes shown in boxed region of Fig. 1A following vgRNA transfection and T3A infection is shown. Black dots denote genes that are not expressed significantly differently in the two treatments. Red dots represent genes that are expressed to a significantly lower extent in T3A infected cells. iRegulon analyses of both sets of genes is also shown. (D, E) ATCC L929 cells were adsorbed with PBS (mock) or 10 PFU/cell of T3A. Following incubation at 37°C for 20 h, cells were transfected with 0.5 μg viral RNA for 7 h. Total RNA was extracted and subjected to RNA-seq analyses. (D) A heat map comparing expression of genes shown in boxed region of Fig. 1A following vgRNA transfection of mock infected and T3A infected cells is shown. (E) A scatterplot comparing expression of genes shown in boxed region of Fig. 1A following vgRNA transfection of mock infected and T3A infected cells is shown. Black dots denote genes that are not expressed significantly differently in the two treatments. Red dots represent genes that are expressed to a significantly lower extent in T3A infected cells transfected with vgRNA. iRegulon analyses of both sets of genes is also shown.

To distinguish between these possibilities, we determined whether reovirus infection inhibits vgRNA-induced NF-κB activation. Toward this end, we compared if gene expression in uninfected cells transfected with vgRNA differed from infected cells transfected with vgRNA. As described above, vgRNA transfection of uninfected cells induces expression of 978 genes (Fig. 1A). In infected cells, however, vgRNA failed to induce 13% of these genes (133 of 978) (Fig. 1D, 1E). We used iRegulon to predict that the most likely transcriptional regulator of genes whose expression was inhibited by reovirus infection was NF-κB, with an NES of 6.1. These data allow us to conclude that reovirus blocks NF-κB dependent gene expression even in the presence of a potent agonist.

vgRNA and reovirus infection activate NF-κB downstream via a common set of sensors that detect RNA (15–17, 19). To determine if the inhibitory effect of reovirus on NF-κB dependent gene expression is only restricted to viral RNA-induced gene expression, we used TNFα, a potent stimulator of NF-κB signaling. RNA-seq analyses of uninfected cells treated with TNFα, using the same criteria described above, led to the upregulation of 32 transcripts (Fig. 2A). In contrast, TNFα has no significant effect on gene expression in cells infected with T3A (Fig. 2B, 2C, 2D). These data indicate that infection of cells with T3A results in blockade of NF-κB-dependent transcription. vgRNA and TNFα initiate NF-κB signaling via distinct routes. Therefore, our analyses suggest that reovirus blocks NF-κB signaling at a step that is shared by both signaling pathways.

**Fig 2.**
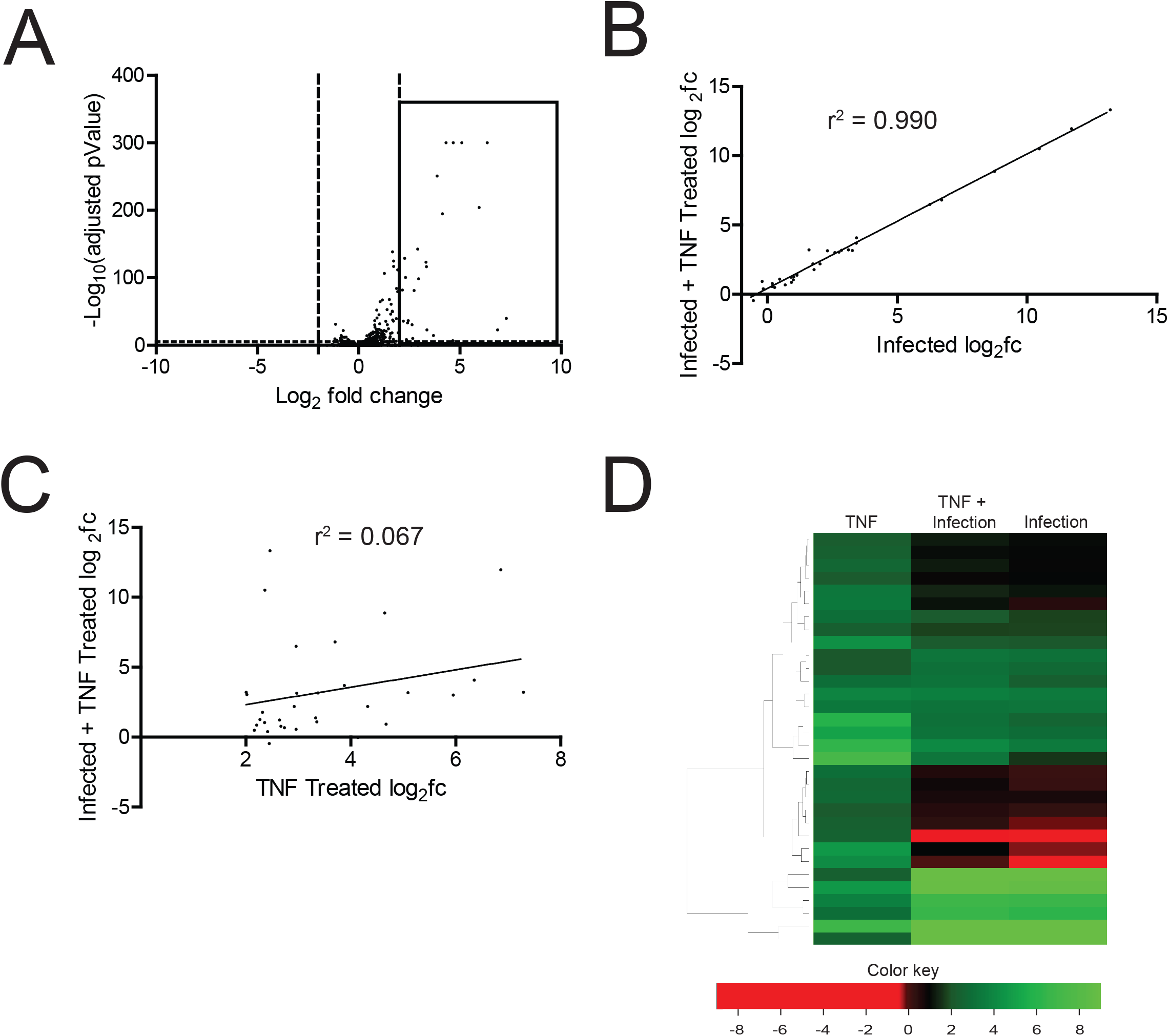
Reovirus strain T3A inhibits TNFα stimulated NF-κB dependent gene expression. (A) ATCC L929 cells were treated with 10 ng/ml TNFα. 1 h following treatment, total RNA was extracted and subjected to RNA-seq analyses. A volcano plot showing genes whose expression is induced > 4-fold (log_2_FC>2) with FDR < 0.05 in comparison to untreated cells are shown within the box. (B, C, D) ATCC L929 cells were adsorbed with 10 PFU/cell of T3A. Following incubation at 37°C for 20 h, cells were treated with 0 or 10 ng/ml TNFα for 1 h. Total RNA was extracted from cells and was subjected to RNA-seq analyses. (B) A scatterplot comparing expression of genes shown in boxed region of Fig. 2A following infection with T3A with or without TNFα is shown. A trendline showing linear regression and coefficient of determination is shown. (C) A scatterplot comparing expression of genes shown in boxed region of Fig. 2A following TNFα treatment of mock infected and T3A infected cells is shown. A trendline showing linear regression and coefficient of determination shown. (D) A heat map comparing expression of genes shown in boxed region of Fig. 2A following TNFα treatment of mock infected and T3A infected cells is shown. Expression of the same set of genes in T3A infected cells is also shown.

### IκB Kinase (IKK) activity is diminished in reovirus-infected cells

To verify our RNA-seq analyses, we measured the capacity of vgRNA and TNFα to induce the expression of an NF-κB target gene in reovirus infected cells using RT-qPCR. For these experiments we monitored the transcript levels of IκBα, an NF-κB target gene. Because the IκBα protein inhibits NF-κB nuclear translocation, its expression serves as a feedback inhibitor of NF-κB activity (20). Consistent with our RNA-seq data, we found that reovirus inhibits IκBα expression to a significant extent following treatment with either agonist (Fig. 3A, 3B). Because the effect of reovirus on both NF-κB agonists was equivalent, we used TNFα for the remainder of our experiments. TNFα treatment of cells should promote nuclear translocation of p65. We measured nuclear p65 levels in mock-infected and reovirus-infected cells treated with TNFα. As expected, TNFα treatment of mock infected cells resulted in an accumulation of p65 in the nucleus within 1 h (Fig. 3C). Prior infection with T3A prevented TNFα driven accumulation of p65 in the nucleus. These data agree with previous evidence indicating that degradation of the IκBα protein is blocked in reovirus infected cells (21). IκBα degradation is initiated by the phosphorylation of IκBα by the IκB Kinase (IKK) complex, which leads to polyubiquitination and subsequent degradation of IκBα by the proteasome (9). Thus, the reduction in nuclear p65 levels in T3A infected cells treated with TNFα may be due to an absence of sufficient levels of active IKK. In addition to IκBα, the IKK complex also phosphorylates p65 at Ser536 prior to nuclear translocation (10). IKK-mediated p65 Ser536 phosphorylation is critical for NF-κB dependent gene expression and is considered to be a marker for IKK activity (10). To determine if IKK activity is compromised in T3A-infected cells, we assessed the capacity of TNFα to promote p65 phosphorylation at Ser536. While TNFα potently induced p65 phosphorylation in mock infected cells, both basal and TNFα induced p65 phosphorylation was dramatically reduced in T3A-infected cells (Fig. 3D). Thus, in reovirus infected cells, p65 nuclear translocation and phosphorylation, both of which require the IKK complex, are inhibited. These data suggest that reovirus may inhibit NF-κB dependent gene expression due to the inactivity of the IKK complex.

**Fig 3.**
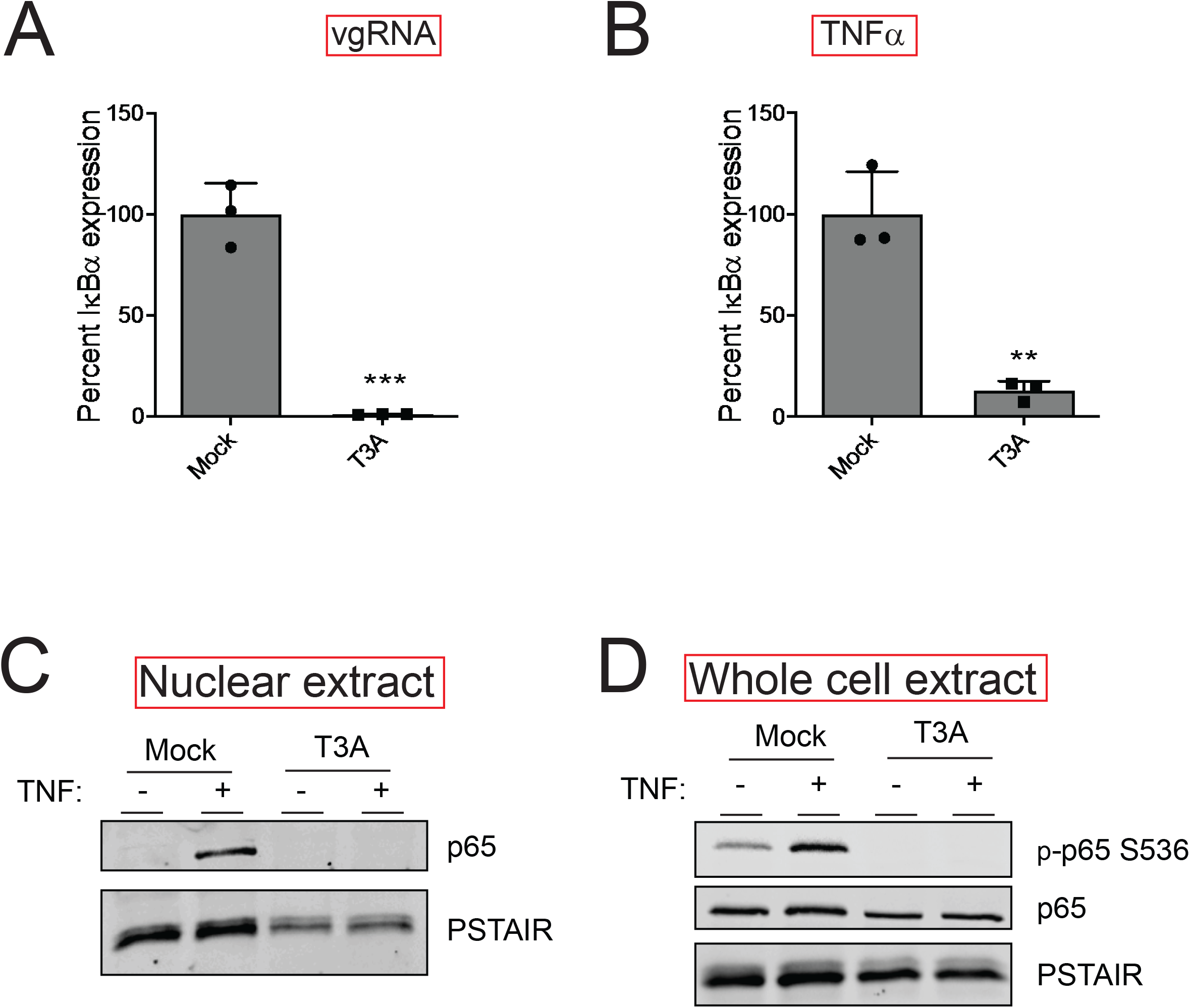
Reovirus inhibits NF-κB signaling upstream of gene expression. (A) ATCC L929 cells were adsorbed with PBS (mock) or 10 PFU/cell of T3A. Following incubation at 37°C for 20 h, cells were transfected with vgRNA and incubated for 7 h. RNA was extracted from cells and levels of IκBα mRNA relative to GAPDH control was measured using RT-qPCR. IκBα expression in mock infected cells treated with agonist vgRNA was set to 100%. Gene expression of each replicate, the mean value, and SD are shown ***, P < 0.001 by Student’s t test in comparison to mock infected cells transfected with vgRNA. (B) ATCC L929 cells were adsorbed with PBS (mock) or 10 PFU/cell of T3A. Following incubation at 37°C for 20 h, cells were treated with 10 ng/ml TNFα and incubated for 1 h. RNA was extracted from cells and levels of IκBα mRNA relative to GAPDH control was measured using RT-qPCR. IκBα expression in mock infected cells treated with agonist TNFα was set to 100%. Gene expression of each replicate, the mean value and SD are shown **, P < 0.01 by Student’s t test in comparison to mock infected cells transfected with vgRNA. (C) ATCC L929 cells were adsorbed with PBS (mock) or 10 PFU/cell of T3A. Following incubation at 37°C for 24 h, cells were treated with 10 ng/ml TNFα and incubated for 1 h. Nuclear extracts were immunoblotted using antiserum specific for p65 or PSTAIR. (D) ATCC L929 cells were adsorbed with PBS (mock) or 10 PFU/cell of T3A. Following incubation at 37°C for 24 h, cells were treated with 20 μM proteasome inhibitor PSI for 1 h, then 10 ng/ml TNFα for 30 min. Whole cell extracts were immunoblotted with antisera specific for p65, p65 Ser536 phosphorylation, and PSTAIR.

### Levels of IKKβ and NEMO are diminished following reovirus infection

To determine the basis of IKK inactivity following infection with T3A, we examined the levels of IKKβ and NEMO, key IKK components that are required for NF-κB activation following TNFα treatment. We found that levels of IKKβ and NEMO are dramatically lower at 12 and 24 h following infection with T3A (Fig. 4A, 4B). In contrast, levels of an upstream signaling protein, RIP1, was unaffected by T3A infection. Similarly, levels of NF-κB constituents p50 and p65 also remained constant. These data indicate that T3A-mediated diminishment in levels of IKKβ and NEMO likely contributes to a reduction in IKK activity and resultant blockade of NF-κB in infected cells. Because IKKβ is the catalytic component of the IKK complex that is required for IκBα and p65 Ser536 phosphorylation, we used IKKβ levels as a surrogate to monitor the mechanism by which IKK activity is diminished following infection.

**Fig 4.**
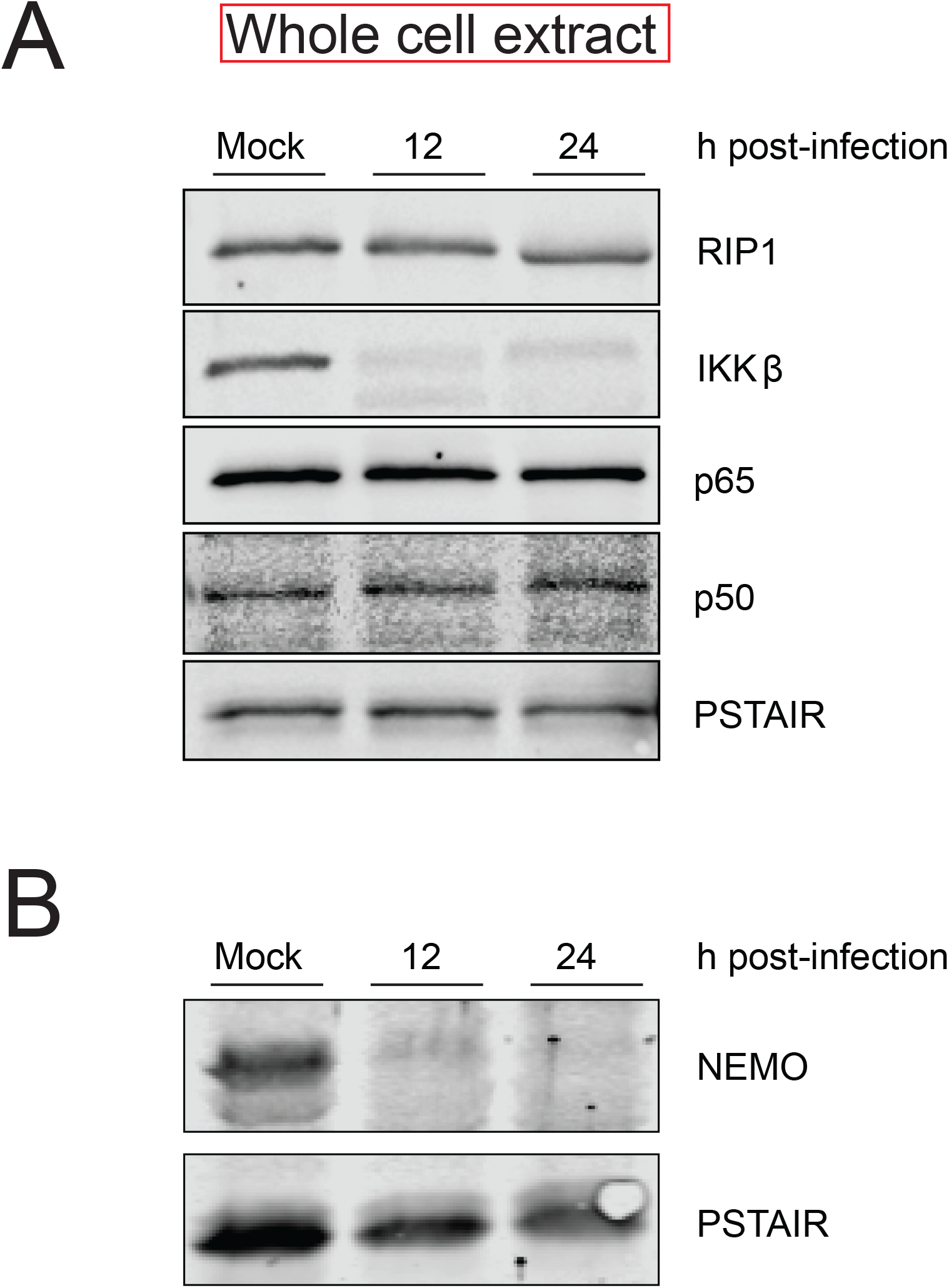
Reovirus infection causes a decrease in IKKβ and NEMO levels. ATCC L929 cells were adsorbed with PBS (mock) or 10 PFU/cell of T3A. Following incubation at 37°C for 12 or 24 h, whole cell extracts were immunoblotted with antisera specific to p50, p65, IKKβ, NEMO, RIP1, and PSTAIR.

### Reovirus gene expression is required for the loss of IKKβ

To determine the stage of the reovirus replication cycle that is required for the loss of the IKK complex and the inhibition of NF-κB, we treated cells with ribavirin, which diminishes viral gene expression (13, 22). We found that ribavirin treatment prevented T3A mediated loss of IKKβ (Fig. 5A). Consistent with this, in cells treated with ribavirin, T3A was no longer able to prevent TNFα-driven nuclear accumulation of p65 (Fig. 5B). Further, ribavirin treatment also reduced the capacity of T3A to block NF-κB dependent gene expression (Fig. 5C). Together these experiments indicate that one or more viral proteins produced following virus infection or a specific event in viral replication triggers the loss of IKKβ.

**Fig 5.**
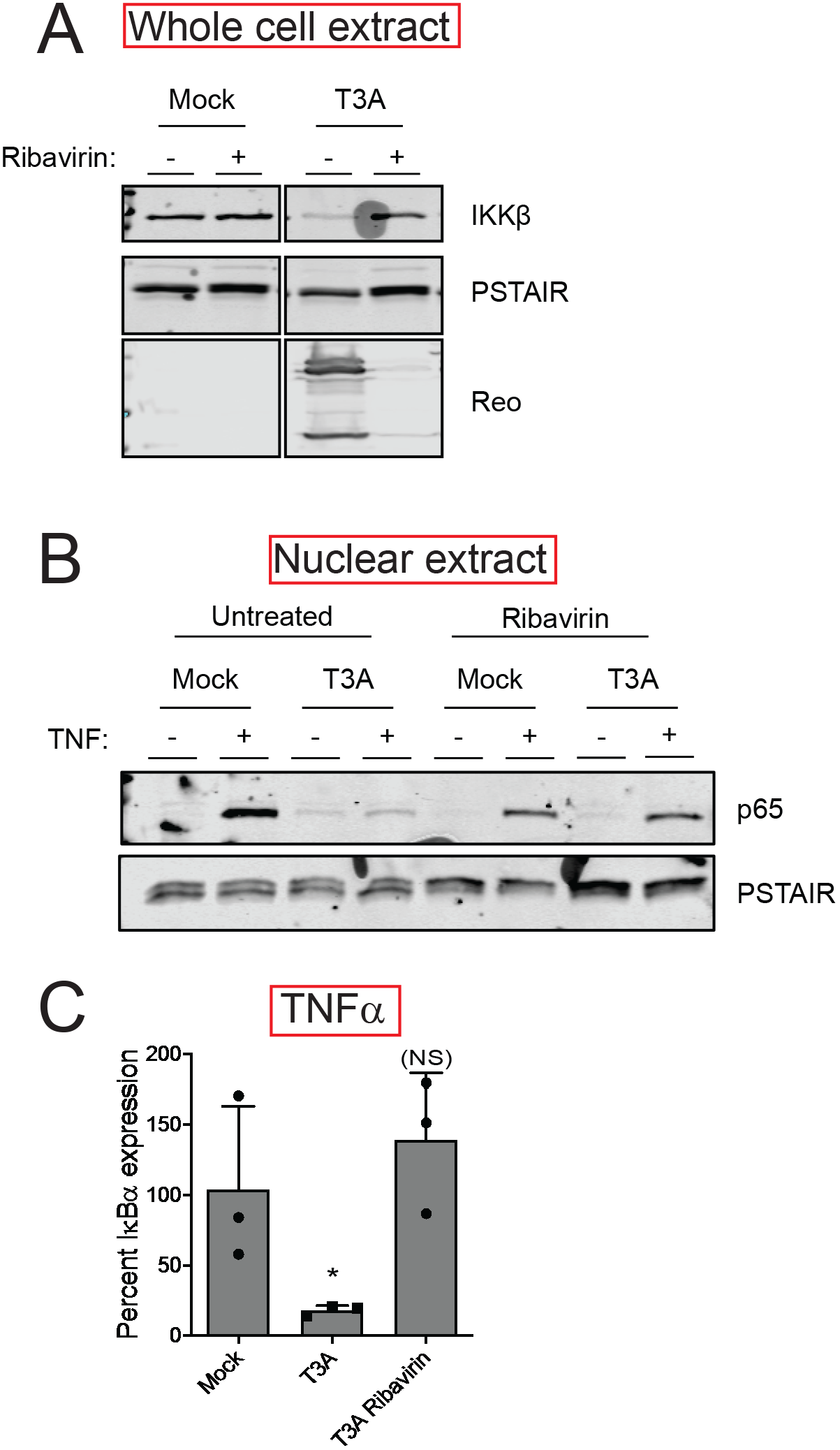
Reovirus gene expression is required for loss of IKKβ. (A) ATCC L929 cells were adsorbed with PBS (mock) or 10 PFU/cell of T3A in the presence of 0 or 200 μM ribavirin. Following incubation at 37^°^C for 24 h, whole cell extracts were immunoblotted with antiserum specific for IKKβ, PSTAIR, and reovirus. (B) ATCC L929 cells were adsorbed with PBS (mock) or 10 PFU/cell of T3A in the presence of 0 or 200 μM ribavirin. Following incubation at 37°C for 24 h, cells were treated with 10 ng/ml TNFα and incubated for 1 h. Nuclear extracts were immunoblotted using antiserum specific for p65 or PSTAIR. (C) ATCC L929 cells were adsorbed with PBS (mock) or 10 PFU/cell of T3A in the presence of 0 or 200 μM ribavirin. Following incubation at 37°C for 24 h, cells were treated with 10 ng/ml TNFα and incubated for 1 h. RNA was extracted from cells and IκBα gene expression was measured using RT-qPCR. Gene expression in mock infected cells treated with TNFα was set to 100%. Gene expression of each replicate, the mean value and SD are shown. *, P < 0.05 by Student’s t test in comparison to mock infected cells treated with TNFα. NS, not significant in comparison to mock infected cells treated with TNFα.

### IKK overexpression restores NF-κB signaling in reovirus infected cells

To define whether blockade of NF-κB by T3A also occurred in other cell lines, we measured the capacity of T3A to influence TNFα induced expression of IκBα in HEK293 cells using RT-qPCR. Analogous to our observation in L929 cells, TNFα failed to induce IκBα gene expression in T3A infected HEK293 cells (Fig. 6A). Phosphorylation of p65 at Ser536 (Fig. 6B) and its nuclear translocation following TNFα treatment were also inhibited (Fig. 6C). Additionally, we noted a decrease in IKKβ levels in comparison to mock infected cells (Fig. 6B). To determine if ectopic overexpression of the IKK complex restores NF-κB signaling in reovirus infected cells, we transfected cells with constructs expressing tagged forms of IKKβ and NEMO. Overexpression of these constructs was sufficient to induce NF-κB signaling in mock infected cells, as marked by p65 Ser536 phosphorylation (Fig. 6D). We found that upon infection of cells with T3A, no decrease in IKKβ levels was observed. Correspondingly, IKK overexpression-induced p65 phosphorylation remained unaffected by reovirus infection. These data further indicate a link between IKKβ levels and NF-κB activity in reovirus infected cells. Thus, we conclude that diminishment in IKKβ and NEMO levels is a key mechanism of reovirus-induced blockade of NF-κB signaling.

**Fig 6.**
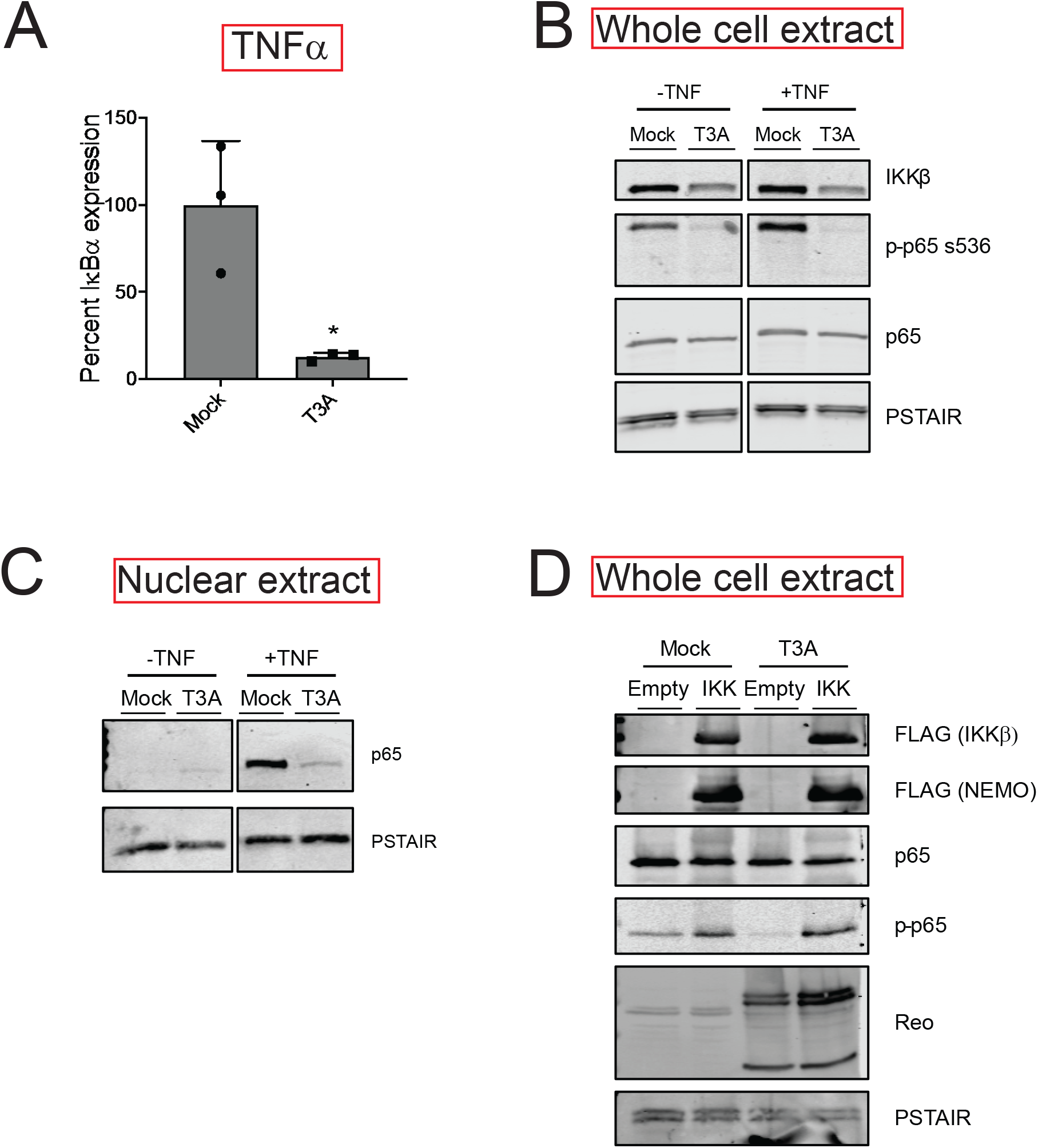
IKK overexpression overcomes reovirus mediated blockade of NF-κB signaling. (A) HEK293 cells were adsorbed with PBS (mock) or 10 PFU/cell of T3A. Following incubation at 37°C for 24 h, cells were treated with 10 ng/ml TNFα and incubated for 1 h. RNA was extracted from cells and IκBα gene expression was measured using RT-qPCR. Gene expression in mock infected cells treated with TNFα was set to 100%. Gene expression of each replicate, the mean value and SD are shown. *, P < 0.05 by Student’s t test in comparison to mock infected cells treated with TNFα. (B) HEK293 cells were adsorbed with PBS (mock) or 10 PFU/cell of T3A. Following incubation at 37°C for 24 h, cells were treated with proteasome inhibitor PSI for 1 h, then 10 ng/ml TNFα for 30 min. Whole cell extracts were immunoblotted using antiserum specific for IKKβ, p65 Ser536 phosphorylation, p65, and PSTAIR. (C) HEK293 cells were adsorbed with PBS (mock) or 10 PFU/cell of T3A. Following incubation at 37°C for 24 h, cells were treated with 10 ng/ml TNFα and incubated for 1 h. Nuclear extracts were immunoblotted using antiserum specific for p65 or PSTAIR. (D) HEK293 cells were transfected with vectors expressing Flag-tagged IKKβ and NEMO. Following incubation at 37°C for 24 h, HEK293 cells were adsorbed with PBS (mock) or 10 PFU/cell of T3A. Following an additional incubation at 37°C for 24 h, whole cell extracts were immunoblotted using antiserum specific for FLAG, p65, p65 Ser536 phosphorylation, reovirus and PSTAIR.

## DISCUSSION

In this study, we sought to evaluate if reovirus infection alters the cellular response to pathogen invasion. Using RNA-seq, which allowed us to examine changes in the global transcriptional landscape, we found that target genes of IRF were induced to a similar extent in cells infected with reovirus and cells transfected with vgRNA. NF-κB target genes, however, were expressed to a much lower extent in infected cells in comparison to cells transfected with vgRNA. Moreover, exogenous NF-κB agonists failed to induce a NF-κB dependent gene expression program in infected cells. These data indicate that NF-κB activity is blocked in infected cells. We found that this blockade of NF-κB dependent gene expression is caused by a loss of IKKβ and NEMO, two critical components of the IKK complex. We propose that reovirus inhibits NF-κB to counter its antiviral effects and produce a cellular environment that is conducive for replication.

We show here that reovirus infection inhibits NF-κB signaling (Fig. 1, 2, 3). In contrast, previous studies demonstrate that reovirus infection leads to the activation of NF-κB (23). While canonical NF-κB signaling requires IKKβ and NEMO, reovirus induced NF-κB activation requires an unusual combination of IKKα and NEMO (24). Reovirus-induced activation of NF-κB occurs early following infection. Recent studies demonstrating a requirement for mitochondrial antiviral-signaling protein (MAVS) for NF-κB activation indicate that vgRNA initiates this response (17). While other work suggests roles for reovirus μ1 and μ2 proteins in activating NF-κB, it is unclear if this effect is through controlling the exposure of vgRNA or via another mechanism (25–27). Regardless, NF-κB activation early in infection does not require viral gene expression. In contrast, our studies presented here indicate that viral gene expression is required for blockade of NF-κB (Fig. 5). Thus, detection of viral RNA activates NF-κB early in infection and expression of one or more viral gene products following establishment of infection results in blockade NF-κB, limiting further signaling through this pathway. Biphasic regulation of NF-κB was also previously suggested (21, 28). Our work presented here provides an explanation for this phenomenon. Serotype-specific capacity of reovirus to inhibit NF-κB is genetically linked to the genome segment encoding the reovirus attachment protein σ1 (28). Because σ1 properties impact the efficiency of infection and the level of viral gene expression (29), we think that the genetic link between σ1 and NF-κB is indirect and the viral factor responsible for diminishment of IKK levels and blockade of NF-κB signaling remains unknown.

NF-κB is an effective pro-inflammatory signaling pathway that curbs infection. Pathogens therefore have evolved mechanisms to limit its activity. Infection with human coronavirus causes a loss of both IKKβ and NEMO through an unknown mechanism (30). The MCMV protein M45 targets NEMO for autophagolysosomal degradation (31). Shigella, an intracellular pathogenic bacteria secretes an effector with E3 ligase activity which targets NEMO for proteasomal degradation (32). In addition to degradation, pathogens also sequester the IKK complex (IAV NS1 protein) or prevent its activation (HCMV, enterovirus)(33–35). Here we show that reovirus infection leads to the loss of both IKKβ and NEMO (Fig. 4). This loss was not due to differences in the steady state levels of IKKβ mRNA in reovirus infected cells (not shown). It was also independent of the effect of reovirus infection on host translation suggesting that IKKβ levels were controlled post translationally (not shown). Reovirus does not encode a protease, ruling out a direct effect of a viral protease on IKKβ. Preventing acid-dependent protease activity also did not restore IKKβ levels, indicating that lysosomal or autophagic degradation does not contribute to IKK loss (not shown). Thus, our work is in contrast to a previous study which suggested that TRIM29 in alveolar macrophages turns over NEMO via lysosomal degradation (36). Blockade of proteasome activity diminished reovirus infection, precluding us from evaluating the role of the proteasome in IKKβ loss following infection (not shown). Thus reovirus infection leads to IKK loss through a post-translational mechanism, likely degradation via a non-lysosomal pathway.

What is the physiologic relevance of blockade of the NF-κB signaling pathway by reovirus? A likely reason is because NF-κB can limit virus replication. Which NF-κB target(s) control virus infection has not been identified. An obvious NF-κB target that could inhibit reovirus infection is IFN. However, consistent with previous work suggesting that certain mouse cell types do not require NF-κB for IFN production (37, 38), we found in our RNA-seq analyses that inhibition of NF-κB did not affect IFN production (Fig. 1). Thus the antiviral effect of NF-κB is independent from IFN. In contrast with IFN, we found that the expression of several other chemokines and cytokines was inhibited in reovirus infected cells. We hypothesize that one or more of these factors negatively regulates reovirus replication.

Two previous studies have suggested that reovirus limits the innate immune response. Reovirus can inhibit IFN production by sequestering IRF3 into viral factories (39). Additionally, reovirus can inhibit IFN signaling by nuclear sequestration of IRF9, which functions with STAT1 and 2 to promote expression of IFN stimulated genes (40). While our work did not directly test these ideas, our gene expression analyses indicate that reovirus does not inhibit the function of IRF3 (Fig. 1). We also do not observe inhibition of the transcriptional complex that contains IRF9 (not shown). Because these previous studies were performed in different cell types and used different reovirus strains, we propose that reovirus has evolved multiple mechanisms to dampen the innate immune response. Our study presented here unveils one such mechanism.

## MATERIALS AND METHODS

### Cells and viruses

Murine L929 cells (ATCC CCL-1) were maintained in Eagle’s minimal essential medium (MEM) (Lonza) supplemented with 10% fetal bovine serum (FBS) and 2 mM L-glutamine. Spinner-adapted L929 cells (obtained from T. Dermody’s laboratory) were maintained in Joklik’s MEM (Lonza) supplemented to contain 5% FBS, 2 mM L-glutamine, 100 U/ml of penicillin, 100 μg/ml of streptomycin, and 25 ng/ml of amphotericin B. HEK293 cells (obtained from M. Marketon’s laboratory) were maintained in Dulbecco’s modified essential medium (DMEM) (Lonza) supplemented with 10% fetal bovine serum (FBS) and 2 mM L-glutamine. Spinner-adapted L929 cells were used for cultivating and purifying viruses and for plaque assays. ATCC L929 cells and HEK293 cells were used for all experiments to assess cell signaling. No differences were observed in permissivity between ATCC L929 cells and spinner-adapted L929 cells. A laboratory stock of T3A (obtained from T. Dermody’s laboratory) was used for infections. Infectious viral particles were purified by Vertrel XF extraction and CsCl gradient centrifugation (41). Viral titer was determined by a plaque assay using spinner-adapted L929 cell with chymotrypsin in the agar overlay.

### Antibodies and reagents

Polyclonal antisera raised against T3D, T1L that have been described (42) were used to detect viral proteins in T3A infected cells. Rabbit antisera specific for IKKβ and p65 Ser536 phosphorylation specific antibody were purchased from Cell Signaling (catalog # 8943, 3033), rabbit antisera specific for p65 and NEMO were purchased from Santa Cruz Biotechnology (catalog # sc-372, sc-8330). Mouse antisera specific for PSTAIR and FLAG was purchased from Sigma-Aldrich (catalog# P7962, F-3165), mouse antisera specific for RIP1 was purchased from BD Biosciences (catalog# 610458) Alexa Fluor-conjugated anti-mouse IgG and anti-rabbit IgG secondary antibodies were purchased from LI-COR. TNFα was purchased from Sigma and used at a concentration of 10 ng/ml. PSI proteasome inhibitor was purchased from Millipore and used at a concentration of 20 μM (catalog# 53-916). Ribavirin was purchased from Sigma-Aldrich and used at a concentration of 200 μM (catalog# R9644)

### Infections

Confluent monolayers of ATCC L929 or HEK293 cells were adsorbed with either PBS or reovirus at the indicated MOI at room temperature for 1 h, followed by incubation with medium at 37°C for the indicated time interval. All inhibitors were added to cells in medium after the 1 h adsorption period.

### Analysis of host gene expression by RNA-seq

Total RNA extracted using Aurum Total RNA Mini Kit (Bio-Rad) was submitted to Indiana University’s Center for Genomics and Bioinformatics for cDNA library construction using a TruSeq Stranded mRNA LT Sample Prep Kit (Illumina) following the manufacturer’s protocol. Sequencing was performed using an Illumina NextSeq500 platform with 75 bp sequencing module generating 38bp paired-end reads. After the sequencing run, demultiplexing with performed with bcl2fastq v2.20.0.422. Sequenced reads were adapter trimmed and quality filtered using Trimmomatic ver. 0.33 (43) with the cutoff threshold for average base quality score set at 20 over a sliding window of 3 bases. Reads shorter than 20 bases post-trimming were excluded (LEADING:20 TRAILING:20 SLIDINGWINDOW:3:20 MINLEN:20). Cleaned reads mapped to GRCm38.p6 mouse genome reference using STAR version STAR_2.5.2b (44). Read pairs aligning to each gene from gencode vM17 annotation were counted with strand specificity using featureCounts tool from subread package (45). The differential expression analysis was performed using DESeq2 version 1.12.3 (46).

iRegulon, a plugin to cytoscape version 3.7.1 was used to predict transcription factor activity based on a differentially expressed gene set (18). A maximum FDR value of motif similarity was set to 0.001. We used a NES value of 3.0 as the minimum cutoff for transcription factor enrichment. Δ NES was calculated as follows: (NES for vgRNA>Mock) – (NES for experimental condition).

### RT-qPCR

RNA was extracted from infected cells, at various times after infection, using Aurum Total RNA Mini Kit (Bio-Rad). For RT-qPCR, 0.5 to 2 μg of RNA was reverse transcribed with the high-capacity cDNA RT kit (Applied Biosystems), using random hexamers. cDNA was subjected to PCR using SYBR Select Master Mix using gene specific primers (Applied Biosystems). Fold increases in gene expression with respect to control samples (indicated in each figure legend) were measured using the ΔΔ*C*_*T*_ method (47). Calculations for determining ΔΔ*C*_*T*_ values and relative levels of gene expression were performed as follows: fold increase in cellular gene expression (with respect to glyceraldehyde-3-phosphate dehydrogenase [GAPDH] levels) = 2^−[(IκBα *CT* − GAPDH *CT*)TNFα − (gene of interest *CT* − GAPDH *CT*control]^

### Preparation of cellular extracts

For preparation of whole-cell lysates, cells were washed in phosphate-buffered saline (PBS) and lysed with 1× RIPA (50 mM Tris [pH 7.5], 50 mM NaCl, 1% TX-100, 1% deoxycholate, 0.1% SDS, and 1 mM EDTA) containing a protease inhibitor cocktail (Roche), 500 μM dithiothreitol (DTT), and 500 μM phenylmethylsulfonyl fluoride (PMSF), followed by centrifugation at 15,000 × *g* at 4°C for 15 min to remove debris. Nuclear extracts were prepared by lysing cells in a hypotonic lysis buffer (10 mM HEPES, 10 mM KCl, 1.5 mM MgCl, 0.5 mM DTT, and 0.5 mM PMSF for 15 min, subsequent addition of 0.5% NP-40, and 10 seconds of vortexing. After centrifugation at 10,000 × g at 4°C for 10 min, nuclear pellet was washed with hypotonic lysis buffer and then resuspended in high-salt nuclear extraction buffer (25% glycerol, 20mM HEPEs, 0.42 M NaCl, 10 mM KCl, 1.5 mM MgCl, 0.5 mM DTT, 0.5 mM PMSF) at 4°C for 1 h. Nuclear extracts were obtained following removal of the insoluble fraction by centrifugation at 12,000 × g at 4°C for 10 min.

### Plasmid transfections

Nearly confluent monolayers of HEK293 cells in 12 well plates were transfected with either 0.5 μg of empty vector or 0.25 μg each of FLAG-IKKβ or FLAG-NEMO expression vector using 1.5 μl Lipofectamine 2000 according to the manufacturer’s instructions. Transfected cells were incubated at 37°C for 24 h prior to infection to allow expression from the plasmids.

### Immunoblot Assay

Protein concentrations were estimated using a DC Protein Assay from Bio-Rad. Equal protein was loaded, and the cell lysates or extracts were resolved by electrophoresis in 10% polyacrylamide gels and transferred to nitrocellulose membranes. Membranes were blocked for at least 1 h in blocking buffer (StartingBlock T20 TBS Blocking Buffer) and incubated with antisera against p65 (1:1,000), p65 p-Ser536 (1:1,000), p50 (1:500), NEMO (1:500), IKKβ (1:1,000), RIP1 (1:1,000), reovirus (1:5,000), FLAG (1:1,000), and PSTAIR (1:5,000) at 4°C overnight. Membranes were washed three times for 5 min each with washing buffer (Tris-buffered saline [TBS] containing 0.1% Tween-20) and incubated with a 1:20,000 dilution of Alexa Fluor-conjugated goat anti-rabbit Ig (for p65, p50, IKKβ, NEMO and reovirus) or goat anti-mouse Ig (for PSTAIR, RIP1, and FLAG) in blocking buffer. Following three washes, membranes were scanned and quantified using an Odyssey infrared Imager (LI-COR).

## ACKNOWLEDGEMENTS

We thank members of our laboratory and the Indiana University Virology community for helpful suggestions. We are also grateful to scientists in the Indiana University Center for Genomics and Bioinformatics.

Research reported in this publication was supported by funds from the National Institute of Allergy and Infectious Diseases under award number R01AI110637 (to P.D.) and by funds from the Indiana Clinical and Translational Sciences Institute under award Number UL1TR002529 from the National Institutes of Health, National Center for Advancing Translational Sciences, Clinical and Translational Sciences Award. The content is solely the responsibility of the authors and does not necessarily represent the official views of the funders.

